# Activation of the Integrated Stress Response overcomes multidrug resistance in FBXW7-deficient cells

**DOI:** 10.1101/2022.01.17.476566

**Authors:** Laura Sanchez-Burgos, Belén Navarro-González, Santiago García-Martín, Héctor Tejero, Marta Elena Antón, Fátima Al-Shahrour, Oscar Fernandez-Capetillo

## Abstract

*FBXW7* is one of the most frequently mutated tumor suppressors, the deficiency of which has been associated with resistance to some anticancer therapies. Through bioinformatic analyses and genome-wide CRISPR screens, we here reveal that FBXW7 deficiency leads to multidrug resistance (MDR), to a bigger extent than well-established MDR-drivers such as overexpression of the drug-efflux pump ABCB1. Proteomic data from FBXW7-deficient cancer cells identify the upregulation of mitochondrial function as a hallmark of FBXW7 deficiency, which has been previously linked to an increased resistance to chemotherapy. Accordingly, genetic or chemical targeting of mitochondria is preferentially toxic for FBXW7-deficient cells *in vitro* and *in vivo*. Mechanistically, we show that the toxicity associated with therapies that target mitochondrial translation such as the antibiotic tigecycline relates to the activation of the Integrated Stress Response (ISR). Furthermore, while searching for additional drugs that could overcome the MDR of FBXW7-deficient cells, we found that all of them unexpectedly also activated the ISR regardless of their currently accepted mechanism of action. Together, our study reveals that one of the most frequent mutations in cancer reduces the sensitivity to the vast majority of available therapies, and identifies a general principle to overcome such resistance.

## INTRODUCTION

Resistance to therapy has been estimated to contribute to treatment failure in up to 90% of cancer patients and remains one of the fundamental challenges in cancer^1,2^. This is also true in the context of immune therapies, the efficiency of which is limited by mutations that reduce antigen presentation or inflammatory signaling^3^. Accordingly, “to develop ways to overcome cancer’s resistance to therapy” was one of the 10 recommendations made from the Blue Ribbon Pannel associated with the Cancer Moonshot initiative of the National Cancer Institute^4^. Starting in the 1960s^5^, intensive research has led to the identification of multiple mutations involved in the resistance to single-agents, leading to the development of targeted therapies that can overcome the resistance. Examples of this are the discovery of MEK reactivation in RAF inhibitor-treated BRAF-mutant melanoma cells, or the emergence of drug-resistant mutations in EGFR in lung cancer, both of which have led to new treatment strategies with improved success^6,7^. Unfortunately, cancer patients under treatment often acquire multidrug resistance (MDR)^8^, which greatly limits their subsequent therapeutic opportunities.

Two of the best known mediators of MDR are the activation of efflux pumps such as ABCB1 that limit intracellular drug concentrations^9^ and the dysregulation of the intrinsic apoptotic pathway through the upregulation of anti-apoptotic proteins like MCL1^10^. Besides the contribution of specific genetic determinants, several evidences indicate that phenotypic changes can also modify the response to therapy in cancer cells. Examples of this include the epithelial-to-mesenchymal transition (EMT)^11^, transdifferentiation into another cell type^12^ or entering into a diapause-like state^13^, all of which have been shown to lead to an increased resistance to chemotherapy. In this regard, an enhanced mitochondrial activity has recently been associated with the resistance to several individual agents^14–23^ as well as to MDR^24–26^. Accordingly, targeting mitochondrial function has emerged as an interesting therapeutic opportunity to overcome drug resistance^27,28^. Furthermore, some tumors such as acute myeloid leukemia (AML)^29^, B-RAF driven melanomas^26^ or C-MYC driven lymphomas^30^ are specifically dependent on mitochondrial translation, which renders them sensitive to certain antibiotics such as tigecycline that affect the function of eukaryotic ribosomes due to their structural resemblance to those from bacteria^31,32^.

FBXW7 is the substrate receptor component of the Skp1-Cdc53/Cullin-F-box-protein (SCF) ubiquitin ligase complex, which mediates the degradation of important oncoproteins such as Cyclin E1 (CCNE1)^33^, MYC^34^, JUN^35^ and NOTCH1^36^ upon their phosphorylation on CDC4 phosphodegron (CPD) domains. In fact, *FBXW7* is one of the 10 most frequently mutated genes in human cancers^37^, due to either inactivating mutations and/or allelic loss^38,39^. Moreover, mutations in *FBXW7* are amongst the most significantly associated with poor survival across all human cancers^40^. Besides its oncogenic potential, loss of FBXW7 has also been linked to an increased resistance to various chemotherapies^41^ and immunotherapy^42^. Furthermore, forward genetic screens have found an enrichment of *FBXW7* mutations among those that drive resistance to various anticancer agents^43–46^. While early works indicated that the increased resistance of FBXW7-deficient cells to agents such as Taxol was due to the stabilization of the antiapoptotic factor MCL1^47^, other mechanisms such as the induction of an EMT have been also proposed to modulate drug sensitivities in *FBXW7*-mutant tumors^48^. We here systematically addressed the impact of FBXW7 deficiency in the response to anticancer therapies, and identify a general strategy to overcome therapy resistance in *FBXW7-*deficient cancer cells.

## RESULTS

### FBXW7 deficiency leads to multidrug resistance

We previously generated mouse embryonic stem cells (mES) harboring a doxycycline-inducible Cas9 and used them to conduct forward genetic screens in order to identify the mechanisms of resistance to inhibitors of the ATR kinase^49^. Following the same pipeline (described in **Fig. S1A**), we performed genetic screens to identify mutations that confer resistance to various cytotoxic agents such as cisplatin, rigosertib or ultraviolet light (UV). When analyzing the sgRNAs present in clones of mutant ES that had become resistant to these treatments, we noted a high frequency of sgRNAs targeting *Fbxw7* (**Fig. S1B**). A similar enrichment of *Fbxw7*-targeting sgRNAs was observed in pools of treatment-resistant mutagenized ES populations that were analyzed by sequencing (**Fig. S1C**). Given the previous literature linking *FBXW7* mutations to resistance to various cancer therapies, we wondered to what extent these observations were reflecting a more general phenomenon and whether FBXW7 deficiency could lead to MDR.

To test the impact of FBXW7 deficiency in the response to cancer therapies, we generated *Fbxw7* wild type (WT) and knockout mES cell lines by CRISPR editing (**Fig. S2A**) which constitutively expressed the fluorescent proteins EGFP or RUBY3, respectively. We then evaluated the effect of different drugs in co-cultures of *Fbxw7*^+/+^ and *Fbxw7*^-/-^ mES by monitoring the evolution in the percentages of EGFP- and RUBY3-positive cells by FACS. Consistent with the reported resistance of *FBXW7-*mutant human cancer cells to paclitaxel^47,50^, oxaliplatin^51,52^, 5-fluorouracil (5-FU)^53,54^ and doxorubicin^55^, *Fbxw7*^-/-^ mES cells became significantly enriched after 48hrs of culture in the presence of these drugs (**Fig. S2B**). Next, and to systematically address the response of *Fbxw7*-deficient cells to chemotherapy, we used the same *Fbxw7*^+/+^*/Fbxw7*^-/-^ competition assay to evaluate a chemical library of 114 FDA-approved antitumoral compounds. Strikingly, this analysis revealed a strong selection for *Fbxw7*-deficient mES cells upon treatment with many different drugs, which was more pronounced with the compounds that had the highest toxicity (**Fig. 1A,B**).

**Fig. 1.**
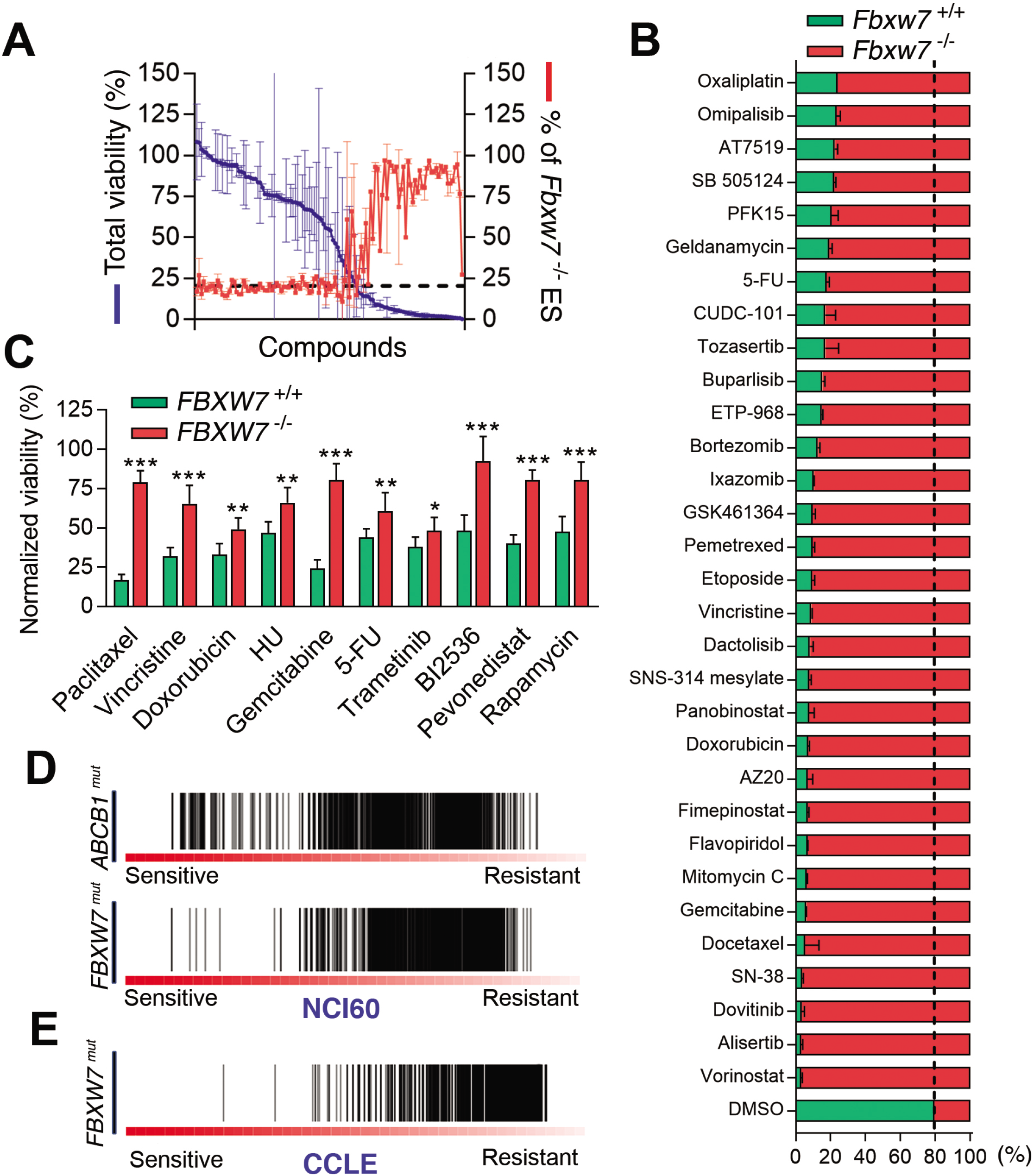
FBXW7 deficiency leads to multidrug resistance. (**A**) Representation of the percentage of total cell viability (left Y-axis; blue) and the percentage of *Fbxw7*^-/-^ mES cells 48h after treatment with 114 different FDA-approved compounds (5 μM). The culture started with a mix of *Fbxw7*^+/+^ and *Fbxw7*^-/-^ mES at a 3:1 ratio. Error bars indicate SD. Cell percentages were quantified by high-throughput flow cytometry. (**B**) Percentages of *Fbxw7*^+/+^ (green) and *Fbxw7*^-/-^ (red) mES cells from the experiment defined in (**A**) with the indicated drugs. (**C**) Percentage of viable *FBXW7*^+/+^ (green) and *FBXW7*^-/-^ (red) DLD-1 cells upon treatment with paclitaxel (40nM), vincristine (10nM), doxorubicin (25nM), hydroxyurea (HU, 75μM), gemcitabine (10nM), Fluorouracil (5-FU, 10μM), trametinib (5μM), BI2536 (PLK1i, 10nM), pevonedistat (200nM) and rapamycin (10μM) for 72h. DMSO was used to normalize viability except for rapamycin, for which ethanol was used as a control. DAPI staining was used to count nuclei by high-throughput microscopy. Error bars indicate SD (n=3). *p<0.05, **p<0.01, ***p<0.001 (t-test). (**D,E**) Profile of drug responses in *FBXW7*^mut^ and *ABCB1*^mut^ cancer cell lines from the NCI60 (**D**) and CCLE (**E**) datasets. Each line represents the response to a specific compound (the higher the score, the less of a response to the compound). No data associated to *ABCB1* mutations was available at the CCLE.

To confirm whether the widespread resistance to chemotherapy was also seen in human cancer cells, we generated *FBXW7*^+/+^ and *FBXW7*^-/-^ clones in the colorectal adenocarcinoma cell line DLD-1 (**Fig. S2C**). We chose colorectal carcinoma as this is the cancer type with the highest frequency of *FBXW7* mutations according to data available at The Cancer Genome Atlas (TCGA). Similar to our observations in mES, *FBXW7*^-/-^ DLD-1 cells were significantly resistant to 10 anticancer drugs with different mechanisms of action (**Fig. 1C**). To obtain a more general view of how FBXW7 deficiency impacts on the response to anticancer drugs, we interrogated data from the NCI-60 repository, where the response of 60 different cancer cell lines to thousands of compounds is available together with genomic and transcriptomic data for each cell line^56^. Consistent with the known role of ABCB1 as a mediator of MDR, *ABCB1* mutant cells were resistant to the majority of the drugs tested in this dataset (**Fig. 1D**). Strikingly, this trend was even more accentuated in *FBXW7*-mutant cell lines. The MDR phenotype associated to *FBXW7* mutations was also observed by interrogating the Cancer Cell Line Encyclopedia (CCLE), which contains data from 1,072 cell lines^57^ (**Fig. 1E**). Similar analyses performed on the Cancer Therapeutics Response Portal (CTRP)^58^ revealed that the increased resistance to chemotherapy not only correlated with *FBXW7* mutations but also with low mRNA expression (**Fig. S2D**), highlighting the potential of using FBXW7 levels as a general biomarker for drug responses. In support of this view, analysis of survival data from the Genomics Data Commons (GDC) portal^59^ revealed that low levels of *FBXW7* expression significantly correlated with poor survival in cancer patients undergoing any type of therapy (**Fig. S2E,F**). Together, these experiments provide compelling evidence indicating that FBXW7 deficiency leads to a profound MDR phenotype in human cancer cells.

### MCL1 and ABCB1 independently contribute to drug resistance in FBXW7-deficient cells

Given that MCL1 is a FBXW7 target that has been previously shown to contribute to the resistance to certain agents such as paclitaxel^47^, we evaluated if this could explain the MDR phenotype of *FBXW7*-mutant cells. To do so, we generated *MCL1* knockouts in *FBXW7*^+/+^ and *FBXW7*^-/-^ DLD-1 cells by CRISPR editing (**Fig. S3A**) and tested their response to the same 10 drugs that we previously observed were less toxic for FBXW7-deficient cells (**Fig. 1C**). While MCL1 deficiency was able to partly overcome the resistance of *FBXW7*^-/-^ DLD-1 cells to 4 of these drugs, including paclitaxel, it had no significant effect in the other six compounds (**Fig. S3B**). Following a similar strategy, we tested whether deletion of the drug-efflux pump ABCB1 could revert the MDR of FBXW7-deficient cells. Once again, while ABCB1 deficiency was able to significantly sensitize *FBXW7*^-/-^ DLD-1 cells to 6 of the drugs, it had no impact on the others (**Fig. S3C,D**). Importantly, neither MCL1 nor ABCB1 loss were able to overcome the resistance of *FBXW7*^-/-^ cells to some of the compounds such as Trametinib, 5-FU or Gemcitabine. Together, these data imply that while specific factors such as ABCB1 or MCL1 might contribute to the resistance of FBXW7-deficient cells to some chemotherapies, another mechanism must account for the more general MDR phenotype found in these cells.

### FBXW7-deficient cells and tumors present an increase in mitochondrial pathways

To investigate whether phenotypic changes could contribute to the MDR of FBXW7-deficient cells, we compared the proteomes of FBXW7 WT and knockout DLD-1 and mES cells. Known targets of FBXW7 such as MYC, DAB2IP or MED13 were upregulated in FBXW7-deficient cells in both cell types (**Fig. 2A**). Besides individual factors, gene set enrichment analyses (GSEA) from proteins that showed increased levels in *FBXW7*^-/-^ cells revealed a significant enrichment in pathways related to mitochondria (**Fig. 2B**), “mitochondrial translation” showing the highest enrichment (**Fig. 2C**). Western blot and immunofluorescence experiments revealed an increase in mitochondrial volume and in levels of complexes from the oxidative phosphorylation (OXPHOS) pathway in *FBXW7*^-/-^ cells (**Fig. S4**). A similar enrichment of mitochondrial pathways was seen in *Fbxw7*^-/-^ mES cells (**Fig. 2D,E**), and comparative analyses confirmed a generalized increase in the levels of mitochondrial proteins in both cell types (**Fig. 2F**). Finally, we used proteomic data available at the CCLE to investigate if equivalent observations were also seen in 388 cancer cell lines. In fact, these analyses confirmed a significant enrichment of mitochondrial proteins in *FBXW7*-mutant cell lines (**Fig. 2G**). In addition, GSEA analyses identified an enrichment of pathways related to mitochondria and oxidative phosphorylation in *FBXW7-*mutant cell lines from the CCLE panel (**Fig. 2H,I**). Together, these data identify that increased mitochondrial activity is a hallmark of *FBXW7-*deficient cells.

**Fig. 2.**
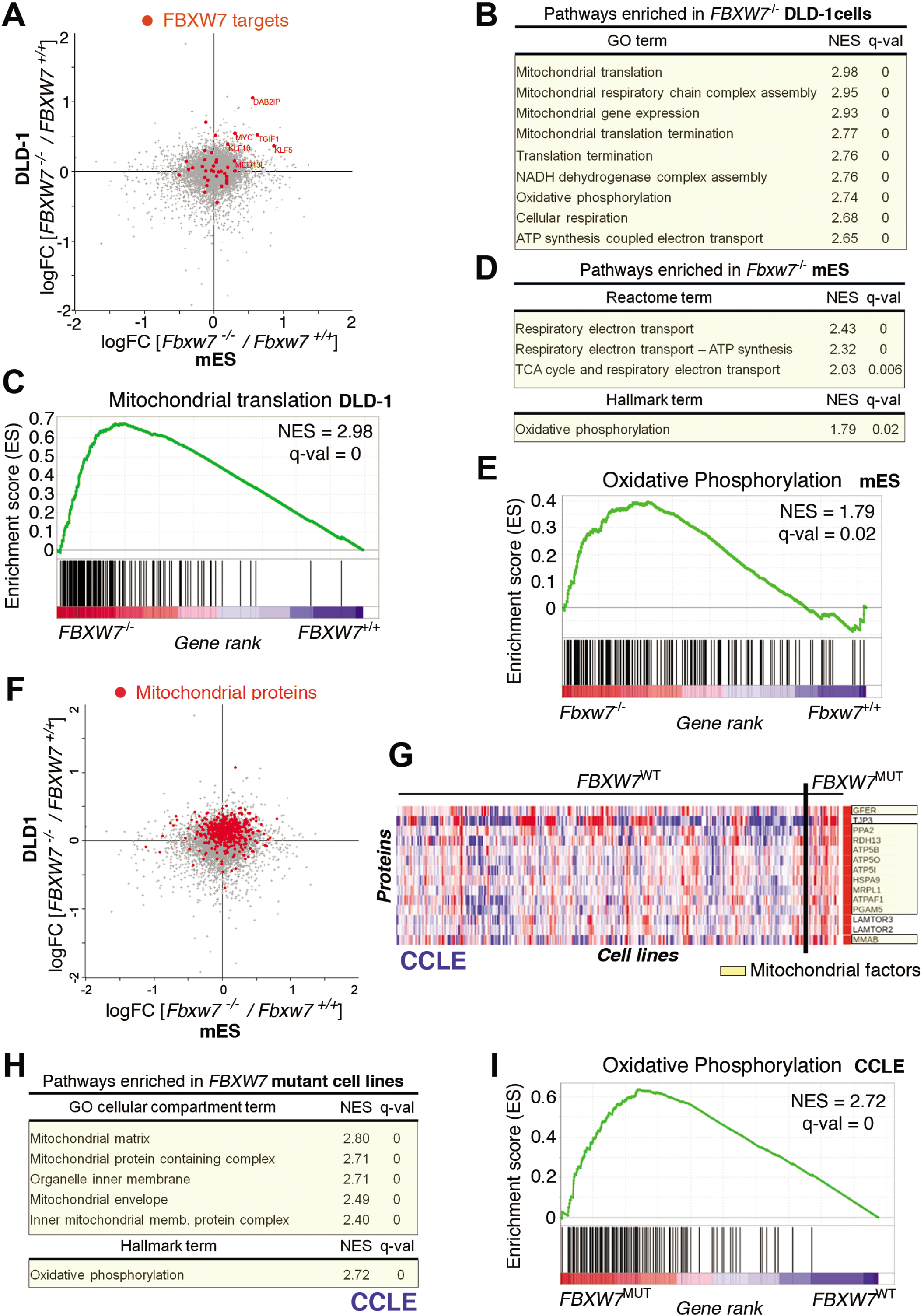
Loss of FBXW7 function is associated to an increased mitochondrial activity. (**A**) Representation of the log_2_FC values from the proteomic analyses of FBXW7 WT and knockout mES (*x*-axis) and DLD-1 (*y-*axis) cells. Known FBXW7 substrates are marked in red. (**B**) GSEA analysis from the proteomic comparison between *FBXW7*^+/+^ and *FBXW7*^-/-^ DLD-1 cells. Normalized enrichment scores (NES), and false discovery rate (FDR) q-values from the most significantly enriched Gene ontology (GO) terms are shown. (**C**) Preranked GSEA on the genes from the ‘Mitochondrial translation’ GO term obtained from the proteomic analysis comparing *FBXW7*^+/+^ and *FBXW7*^-/-^ DLD-1 cells. The heatmap representation illustrates the overall upregulation of these pathways in FBXW7-deficient cells. (**D**) GSEA analysis from the proteomic comparison between *Fbxw7*^+/+^ and *Fbxw7*^-/-^ mES cells. Normalized enrichment scores (NES), and false discovery rate (FDR) q-values from the most significantly enriched Reactome and Hallmark terms are shown. (**E**) Preranked GSEA on the genes from the ‘Oxidative Phosphorylation’ hallmark obtained from the proteomic analysis comparing *Fbxw7*^+/+^ and *Fbxw7*^-/-^ mES cells. (**F**) Representation of the log_2_FC values from the proteomic analyses of FBXW7 WT and knockout mES (*x*-axis) and DLD-1 (*y-*axis) cells (as in (A)). Mitochondrial proteins are marked in red. (**G**) Differential protein expression analysis between *FBXW7*^+/+^ and *FBXW7*^mut^ cancer cell lines from the CCLE. Proteins significantly upregulated in *FBXW7*^mut^ cell lines are displayed, with mitochondrial factors highlighted in yellow. (**H**) GSEA analysis from the proteomic comparison between *FBXW7*^+/+^ and *FBXW7*^mut^ cancer cell lines from the CCLE. Normalized enrichment scores (NES), and false discovery rate (FDR) q-values from the most significantly enriched “GO cellular compartment” and “Hallmark” terms are shown. (**I**) Preranked GSEA on the genes from the ‘Oxidative Phosphorylation hallmark obtained from the proteomic analysis comparing *FBXW7*^+/+^ and *FBXW7*^mut^ cancer cell lines from the CCLE. Note: zero q-values indicate that the value is <10^−4^.

### Targeting mitochondrial function is preferentially toxic for FBXW7-deficient cells

As mentioned above, mitochondrial translation can be targeted by antibiotics that also inhibit the eukaryotic mitochondrial ribosome^31,32^. Competition experiments using *FBXW7*^+/+^ and *FBXW7*^-/-^ DLD-1 cells expressing EGFP and RUBY3, respectively, revealed that several antibiotics led to a depletion of *FBXW7*^-/-^ cells, with tigecycline having the biggest effect (**Fig. 3A,B**). Counting nuclei by High-Throughput Microscopy (HTM) revealed that this effect was due to a preferential toxicity of tigecycline for *FBXW7*^-/-^ DLD-1 cells (**Fig. 3C**). A similar sensitivity to tigecycline was also seen in FBXW7-deficient HeLa and A2780 cells generated by CRISPR, ruling out cell line-specific effects (**Fig. S5**). The preferential toxicity of tigecycline was also confirmed in tumor xenografts of *FBXW7*^-/-^ DLD-1 cells, the growth of which was in contrast unaffected by paclitaxel (**Fig. 3D,E**). Of note, the sensitivity of FBXW7-deficient cells to tigecycline was partly rescued by siRNA-mediated depletion of the oncogene MYC, which is a FBXW7 target previously shown to confer sensitivity to the inhibition of mitochondrial translation^30^ (**Fig. S6A,B**). MYC depletion also reduced the levels of several mitochondrial factors in *FBXW7*^-/-^ DLD-1 cells (**Fig. S6A**). Besides antibiotics, drugs targeting OXPHOS such as IACS-10759 ^60^ or the mitochondrial F1F0 ATPase inhibitor oligomycin were also preferentially toxic for *FBXW7*^-/-^ DLD-1 cells (**Fig. 3F,G**), yet not as much as tigecycline. Finally, and in agreement with the effects observed with chemicals, depletion of the mitochondrial factors TUFM, POLRMT, PTCD3, MRPS27 and UQCRC1 by enzymatically generated short interfering RNAs (esiRNA) was also preferentially toxic for *FBXW7*^-/-^ DLD-1 cells as shown in competition experiments (**Fig. 3H**). Together, these experiments demonstrate that targeting mitochondrial function is able to overcome the widespread resistance to chemotherapy associated to FBXW7 deficiency in cancer cells.

**Fig. 3.**
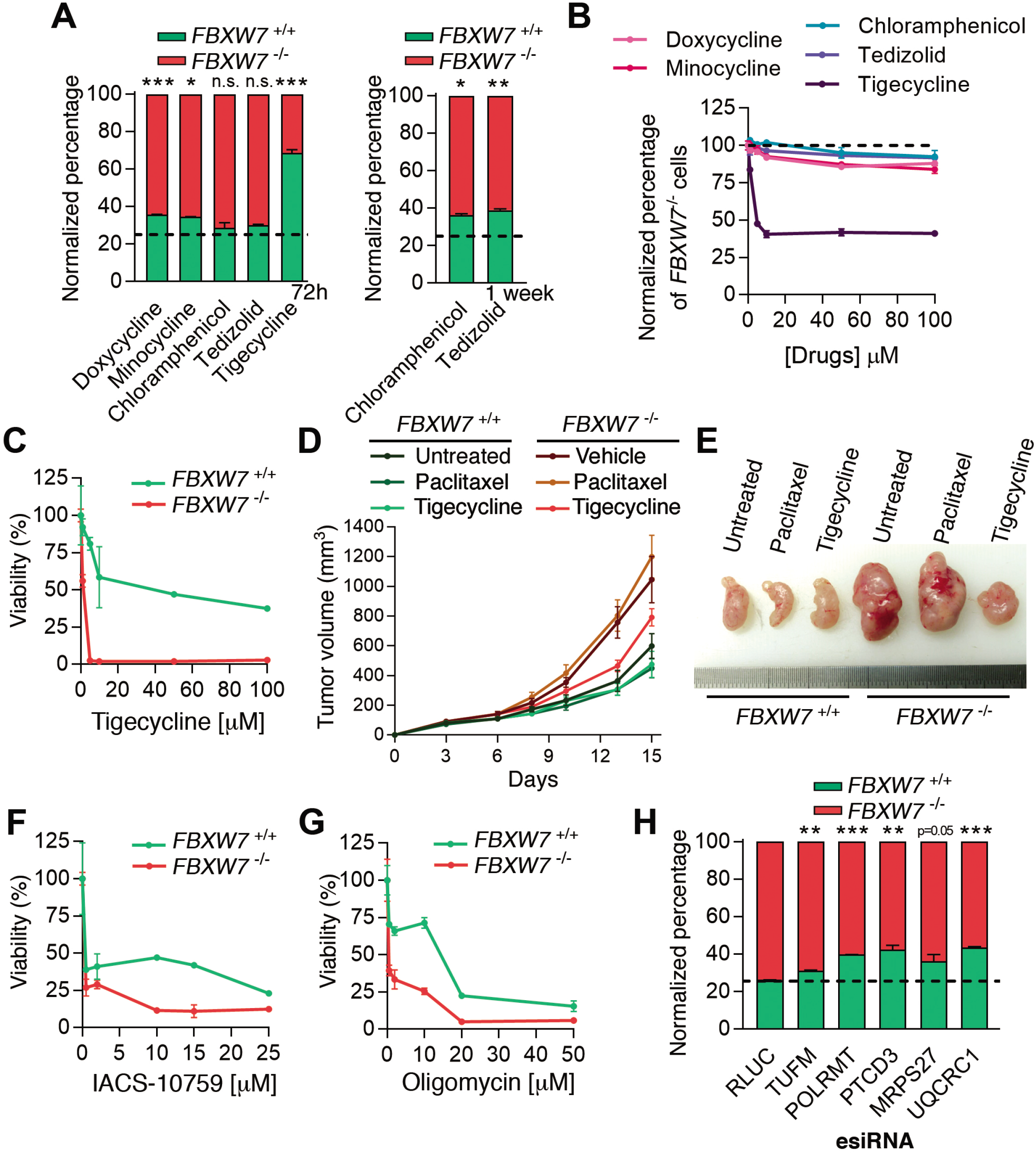
Targeting mitochondrial function is preferentially toxic for FBXW7-deficient cells. (**A**) Percentage of viable *FBXW7*^+/+^ (green) and *FBXW7*^-/-^ (red) DLD-1 cells 72h or 1 week after being treated with the indicated antibiotics. The culture started with a 1:3 ratio of *FBXW7*^+/+^ and *FBXW7*^-/-^ cells. The experiment was repeated three times, and a representative example is shown. Error bars indicate SD. n.s. p>0.05, *p<0.05, **p<0.01, ***p<0.001 (t-test). Cell percentages were quantified by flow cytometry. (**B**) Normalized percentage of *FBXW7*^-/-^ cells using increased doses of the indicated antibiotics following the same experimental approach defined in (**A**). (**C**) Normalized viability of *FBXW7* ^+/+^ and *FBXW7* ^-/-^ DLD-1 cells upon treatment with increasing doses of tigecycline for 72h. Cell nuclei were quantified by high-throughput microscopy (HTM) upon staining with DAPI. The experiment was repeated three times, and a representative example is shown. Error bars indicate SD. (**D**) Tumour growth (in mm^3^) of *FBXW7* ^+/+^ and *FBXW7* ^-/-^ xenografts in nude mice (n=10 animals per group). Treatment with either vehicle, paclitaxel (1,5mg/kg) or tigecycline (50mg/kg) started at day 6 post-tumour-injection, and was administered three times per week. Error bars indicate SEM. (**E**) Representative images of the xenografts defined in (**D**) at day 15. (**F,G**) Normalized viability of *FBXW7*^+/+^ and *FBXW7*^-/-^ DLD-1 cells upon treatment with increasing doses of IACS-10759 (**F**) or oligomycin (**G**) for 72h. Cell nuclei were quantified by HTM as in (**C**). The experiment was repeated three times, and a representative example is shown. Error bars indicate SD. (**H**) Percentage of viable *FBXW7*^+/+^ (green) and *FBXW7*^-/-^ (red) DLD-1 cells 7 days after being transfected with esiRNAs targeting the indicated mitochondrial factors or luciferase (RLUC) as control. The culture started with a 1:3 ratio of *FBXW7*^+/+^ and *FBXW7*^-/-^ cells. The experiment was repeated three times, and a representative example is shown. Error bars indicate SD. **p<0.01, ***p<0.001 (t-test). Cell percentages were quantified by flow cytometry.

### The toxicity of tigecycline is mediated by the integrated stress response

Next, we aimed to understand the mechanism by which tigecycline promotes the preferential killing of FBXW7-deficient cells. To this end, we evaluated the transcriptional changes induced by tigecycline in *FBXW7*^+/+^ and *FBXW7*^-/-^ DLD-1 cells. Consistent with toxicity experiments, tigecycline had a significantly bigger impact on the transcriptome of *FBXW7*^-/-^ cells (**Fig. 4A**). Gene Ontology (GO) analyses revealed that the antibiotic triggered various stress responses, including the endoplasmic reticulum (ER) stress response or the cellular response to arsenate, which were particularly acute in mutant cells (**Fig. S7**). These hallmarks suggested that tigecycline was activating the Integrated Stress Response (ISR), a signaling network that reprograms gene expression to respond to a wide range of insults but which can also promote apoptosis to eliminate the damaged cell^61^. In support of this, tigecycline promoted the nuclear accumulation of the transcription factor ATF4, a hallmark of the ISR (**Fig. 4B**)^62^. Furthermore, tigecycline-induced nuclear translocation of ATF4 was accentuated in *FBXW7*^-/-^ cells, and was reverted by the ISR inhibitor ISRIB^63,64^ (**Fig. 4B**). More importantly, clonogenic survival assays revealed that ISRIB fully rescues the toxicity of tigecycline in both WT and FBXW7-deficient DLD-1 cells, confirming that the cytotoxic effects of the antibiotic are mediated by the ISR (**Fig. 4C**).

**Fig. 4.**
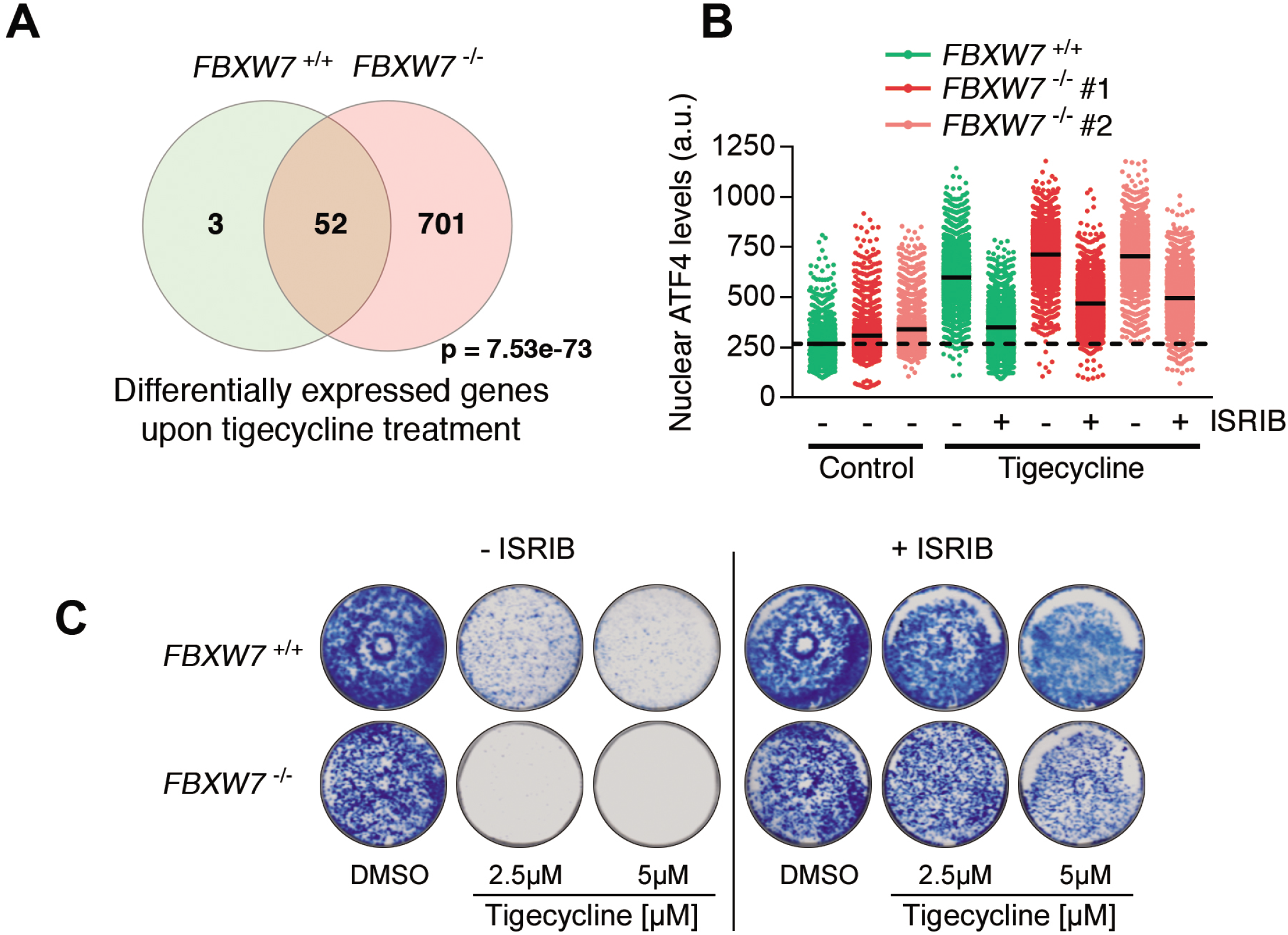
The toxicity of tigecycline is mediated by the ISR. (**A**) Venn Diagram of the genes that are significantly overexpressed upon a 24h treatment with tigecycline (10 μM) in *FBXW7*^+/+^ and *FBXW7*^-/-^ DLD-1 cells (padj < 0.1). Note that the antibiotic has a much wider impact on the mutant cells. The p-value indicates the statistical significance of the overlap between the groups. (**B**) Nuclear ATF4 levels quantified by HTM in *FBXW7*^+/+^ and *FBXW7*^-/-^ DLD-1 cells upon treatment with tigecycline (10 μM) with or without the ISR inhibitor ISRIB (50nM) for 3h. This experiment was performed 3 times, and a representative example is shown. (**D**) Clonogenic assays in *FBXW7*^+/+^ and *FBXW7*^-/-^ DLD-1 cells treated with the indicated doses of tigecycline with or without 50nM ISRIB. Control plates were treated with DMSO. This experiment was performed 3 times, and a representative example is shown.

### Drugs activating the ISR overcome the MDR of FBXW7-deficient cancer cells

Finally, we sought to identify additional drugs that could overcome the widespread resistance to chemotherapy of FBXW7-deficient cells. To do so, we interrogated the Connectivity Map (CMap) dataset, which contains the transcriptional signatures triggered by thousands of drugs in cancer cells^65^, in order to identify drugs that elicit a transcriptional signature similar to that of tigecycline. Consistent with our RNAseq results, the signatures induced by the mitochondrial poison oligomycin and the ISR activators tunicamycin and salubrinal showed the highest similarity to that from tigecycline (**Fig. 5A**). In addition, and like tigecycline or oligomycin, *FBXW7*^-/-^ cells were also sensitive to tunicamycin, confirming that activating the ISR is preferentially toxic for FBXW7-deficient cells (**Fig. 5B**). Interestingly, the list of drugs with signatures most similar to tigecycline included other drugs with seemingly distinct mechanisms of action such as B-RAF inhibitors (PLX-4720 and vemurafenib), broad spectrum tyrosine kinase inhibitors (sorafenib and dasatinib) and EGFR inhibitors (erlotinib and gefitinib). Given the similarity of the transcriptional signatures triggered by these drugs to those of tigecycline and tunicamycin, we wondered if they also activated the ISR. In fact, all of these drugs promoted the nuclear accumulation of ATF4, which was reverted by ISRIB (**Fig. 5C** and **Fig. S8A**). In addition, these drugs promoted the accumulation of CHOP, a transcription factor that mediates the apoptosis triggered by the ISR (**Fig. S8B**). As to whether these drugs were able to overcome the MDR of FBXW7-deficient cells, competition experiments using *FBXW7*^+/+^ and *FBXW7*^-/-^ DLD-1 cells showed a depletion of *FBXW7*^-/-^ cells upon treatment with all of the drugs (**Fig. 5D**). Similarly, HTM-mediated quantification of nuclei confirmed that these compounds were preferentially toxic for *FBXW7*^-/-^ cells, in a manner that could be rescued by ISRIB (**Fig. 5E**). Noteworthy, drugs to which FBXW7-deficient cells are resistant such as paclitaxel or trametinib failed to activate the ISR as measured by the accumulation of ATF4 or CHOP (**Fig. 5C** and **S8B**).

**Fig. 5.**
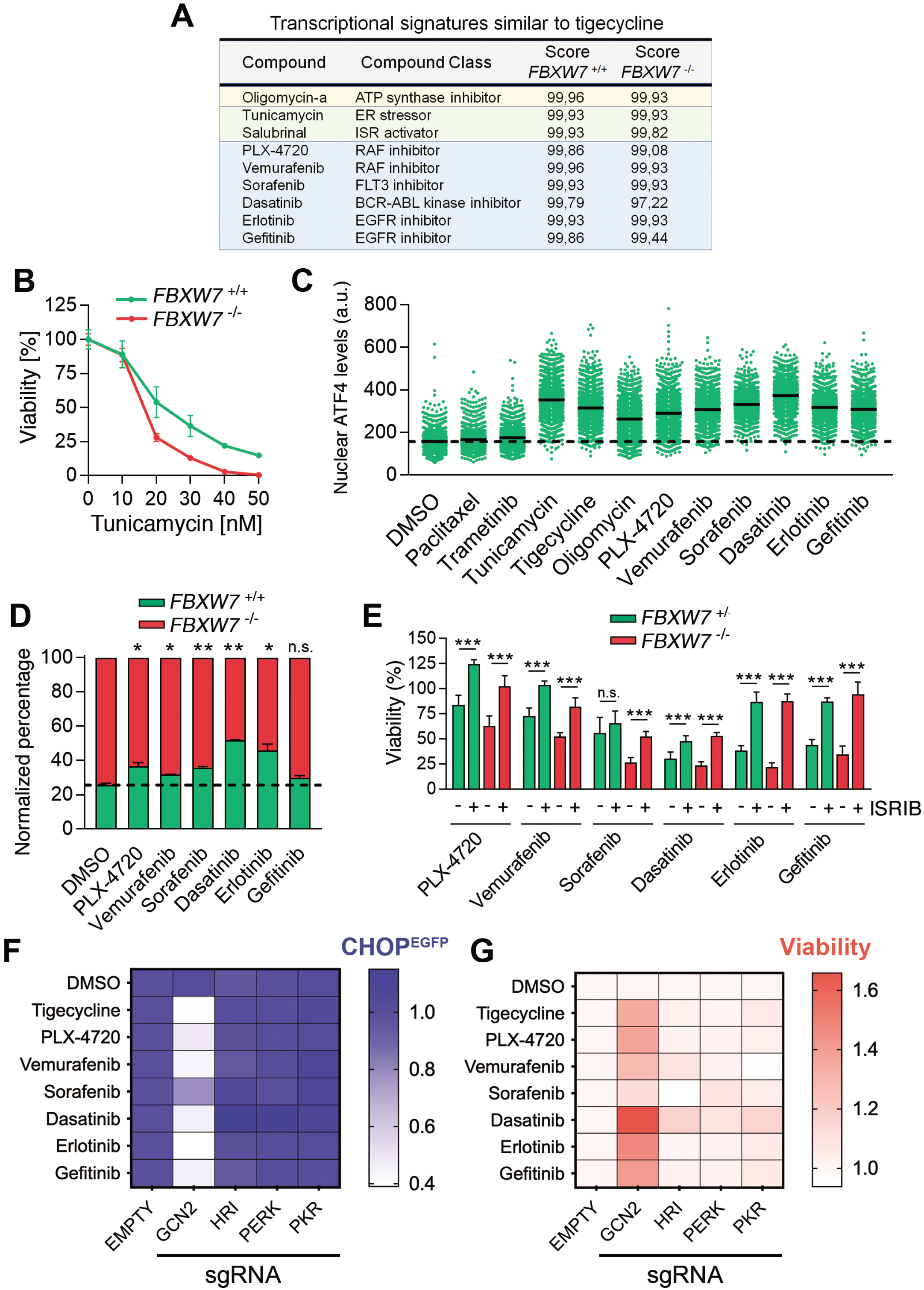
ISR-activating drugs overcome the MDR of FBXW7-deficient cells. (**A**) Compounds with a transcriptional signature similar to that triggered by tigecycline identified by CMap using RNAseq data from tigecycline-treated *FBXW7*^+/+^ and *FBXW7*^-/-^ DLD-1 cells. Compound name, compound class and their similarity scores from both genotypes are shown. Mitochondrial poisons (yellow), ISR-activating compounds (green) and additional compounds (blue) are highlighted. (**B**) Normalized viability of *FBXW7* ^+/+^ and *FBXW7*^-/-^ DLD-1 cells upon treatment with increasing doses of tunicamycin for 72h. Cell nuclei were quantified by high-throughput microscopy (HTM) upon staining with DAPI. The experiment was repeated three times, and a representative example is shown. Error bars indicate SD. (**C**) Nuclear ATF4 levels quantified by HTM in DLD-1 cells upon treatment with the indicated drugs at 10μM (except tunicamycin (1 μM) and paclitaxel (250 nM)) for 3h. This experiment was performed 3 times, and a representative example is shown. (**D**) Percentage of viable *FBXW7*^+/+^ (green) and *FBXW7*^-/-^ (red) DLD-1 cells treatment with the indicated drugs at 15 μM for 72h. The culture started with a 1:3 ratio of *FBXW7*^+/+^ and *FBXW7*^-/-^ cells. DMSO was used as a control. The experiment was repeated three times, and a representative example is shown. Error bars indicate SD. n.s.: non-significant, **p<0.01, ***p<0.001 (t-test). Cell percentages were quantified by flow cytometry. (**E**) Normalized viability of *FBXW7*^+/+^ and *FBXW7*^-/-^ cells upon treatment with 10μM of PLX-4720, vemurafenib, erlotinib and gefitinib, and 5μM of sorafenib and dasatinib for 72h, in the presence or absence of ISRIB (50nM). Cell nuclei were quantified by high-throughput microscopy (HTM) upon staining with DAPI. Error bars indicate SD (n=3). n.s. p>0.05, *p<0.05, **p<0.01, ***p<0.001 (t-test). (**F**) Heatmap representing of the EGFP median fluorescence intensity (MFI) in DLD-1 cells stably transfected with a reporter construct where a destabilized EFGP is under the control of the human *CHOP* promoter, after treatment with the indicated drugs. Cells were infected with lentiviral vectors expressing mCherry and sgRNAs against the four ISR kinases as well or an empty vector. Cell percentages were quantified by flow cytometry. EGFP MFI ratios between mCherry positive and negative cells were first calculated and then normalized to the levels observed in the controls for each treatment. (**G**) Heatmap representing the viability (calculated as the percentage of mCherry positive cells) in DLD-1 cells treated as in (**F**). Percentages were normalized first to the DMSO control values and then to the ones found on the empty vector for each treatment.

Finally, we investigated which one of the 4 kinases that activate the ISR (HRI, GCN2, PERK or PKR) mediated the effects of these drugs on the ISR. To do so, we generated DLD-1 cells carrying a transcriptional reporter where a destabilized EGFP is placed under the control of the human CHOP promoter. These cells were subsequently infected with lentiviral viruses expressing mCherry and sgRNAs targeting ISR kinases or an empty vector. Flow cytometry analyses revealed that GCN2 deletion prevented the expression of CHOP triggered by tigecycline and by the drugs that overcome the MDR of FBXW7-deficient cells (**Fig. 5F**). Furthermore, infection with lentiviruses expressing sgRNAs against GCN2, but not against HRI, PERK or PKR, reduced the toxic effect of all of these drugs (**Fig. 5G**). Collectively, these results identify the activation of the ISR as a general strategy to overcome drug resistance in FBXW7-deficient cancer cells, and suggest that the toxic effects of several anticancer drugs might be partly contributed by a previously unknown effect of the compounds in activating the GCN2-dependent branch of the ISR.

## DISCUSSION

We here show that FBXW7 deficiency, one of the most frequent events in human cancer^37^, increases the resistance to the vast majority of available anticancer therapies, likely contributing to the bad prognosis that is associated to *FBXW7* mutations^40^. Since the increased resistance to chemotherapies is also associated to reduced *FBXW7* levels, this opens the possibility of using FBXW7 expression as a biomarker to predict an unfavorable response to treatment. We further reveal that the MDR is associated to an increased mitochondrial activity, rendering FBXW7-deficient tumors to reagents that target mitochondrial function. Noteworthy, our proteomic analysis is consistent with the analysis of transcriptional signatures from The Cancer Genome Atlas that indicate an increase in mitochondrial gene expression in *FBXW7-*deficient tumors^66^. As for the mechanism behind this phenomenon, several evidences indicate an important role for MYC. First, MYC is an FBXW7 target^34^ that stimulates mitochondrial biogenesis and activity^24,67^. Second, increased levels of MYC have been shown to increase resistance to several therapies^24,68^. Finally, MYC overexpression renders cancer cells sensitive to targeting mitochondrial activity with antibiotics^69^ or drugs targeting OXPHOS^70^. Given the increasing availability of chemical- or peptide-based therapies for targeting MYC^71^, whether these agents might be useful as a general strategy to overcome the MDR in cancer emerges as an interesting possibility.

Targeting mitochondrial activity has been a long-debated approach in cancer therapy. Early clinical efforts faced either toxicities of drugs targeting OXPHOS or limited efficacy from tetracycline antibiotics^27,28^. Nevertheless, there is an intense preclinical development of additional chemotherapies that affect mitochondrial function which include inhibitors of mitochondrial transcription^72^, inhibitors of the METTL8 RNA methyltransferase^73^, and even the use of a diet low in valine, which has been recently shown to preferentially impair mitochondrial function and exert antitumoral properties^74^. Interestingly, a recent study also revealed that mitochondrial activity can be used to identify a glioblastoma subtype that is vulnerable to OXPHOS inhibitors^75^. Despite this renewed interest in targeting mitochondria in cancer, important questions still remain to be addressed. In our opinion, and before these strategies are brought to clinical trials, it would be important to identify the genetic determinants that modulate the response to these treatments, both to select patients that are most likely to respond but also to avoid their use in patients carrying mutations that limit their efficacy.

Finally, our study indicates that the effect of drugs targeting mitochondria in overcoming the MDR of FBXW7-deficient cells is associated with their capacity to activate the ISR. This data is consistent with the fact that mitochondrial stress triggers an ATF4-dependent stress response in mammalian cells^62^. Furthermore, the antitumoral effects of antibiotics were also previously associated with the ISR^76^. Intriguingly, activation of the ISR has also been shown to stimulate mitochondrial translation^26^, so that it is possible that the increased mitochondrial dependence of FBXW7-deficient cancer cells is secondary to an active ISR. Similarly to mitochondrial targeting drugs, therapies that stimulate the ISR are also being explored in cancer therapy^77^. Despite focused efforts to generate selective activators of the ISR, we here report the surprising discovery that several drugs with distinct targets and mechanisms of action are activators of the ISR. This raises the question as to what extent the antitumoral effects of these drugs might be partly due to their effects on the ISR, and leads us to propose that this phenotype should be routinely analyzed when developing new anticancer therapies. Consistent with our findings, a recent study has identified that several ATP-competitive kinase inhibitors directly bind and activate GCN2, thereby activating the ISR^78^. To what extent activating the ISR is a general strategy to overcome MDR regardless of FBXW7 is also an important question to be addressed in future studies.

## ACKNOWLEDGEMENTS

We would want to thank Drs. Daniela Hühn and Andrés López-Contreras for insightful comments on the manuscript, as well as Javier Muñoz and Eduardo Zarzuela for their help with proteomic analyses. Research was funded by grants from the Spanish Ministry of Science, Innovation and Universities (RTI2018-102204-B-I00, co-financed with European FEDER funds) and the Spanish Association Against Cancer (AECC; PROYE20101FERN) to OF; from the Spanish Ministry of Science, Innovation and Universities (RTI2018-097596-B-I00, (AEI/10.13039/501100011033 MCI/FEDER, UE), co-financed with European FEDER funds) to FA; and a PhD fellowship from La Caixa Foundation and the Marie Skłodowska-Curie European Union’s Horizon 2020 actions (LCF/BQ/IN17/11620001) to LS.

## AUTHOR CONTRIBUTIONS

L.S. contributed to most experiments and data analyses and to the preparation of the figures. B.N. contributed to experiments on the mechanism of action of tigecycline. S.G., H.T. and F.A. performed most bioinformatic analyses. M.A. provided help in mouse xenograft studies. O.F. supervised the study and wrote the MS.

## MATERIALS AND METHODS

### Cell culture

All cells were grown at 37°C in a humidified air atmosphere with 5% CO_2_ unless specified. mES were grown on gelatine and feeder layers, using DMEM (high glucose) (Invitrogen) supplemented with 15% knockout serum replacement (Invitrogen), LIF (1000 U/ml), 0.1 mM non-essential amino acids, 1% glutamax, and 55mM β-mercaptoethanol. Wild type mES cells (R1) were obtained from the American Type Culture Collection (ATCC). ES Cas9 clones and loss-of-function libraries were previously generated^49^. Human cancer cell lines HEK-293T, DLD-1 and HeLa cells (ATCC) were cultured in standard DMEM (high glucose) (Sigma, D5796) supplemented with 10% FBS and 1% penicillin/streptomycin. A2780 cells (ATCC) were maintained in RPMI 1640 medium (EuroClone, ECM2001L), 10% FBS and 1% penicillin/streptomycin. For the analysis of *CHOP* transcription, DLD-1 cells were infected with pCLX-CHOP-dGFP lentiviruses (Addgene, 71299), single cell isolated, and selected on the basis of expressing dGFP in response to tunicamycin.

### CRISPR editing

To generate *Fbxw7* knockout ES and *FBXW7*^-/-^ DLD-1, HeLa and A2780 cell lines, cells were independently infected with lentiviral supernatants encoding sgRNAs against *Fbxw7/FBXW7.* Each cell line was also infected with the empty pLentiCRISPR v2 vector (Addgene, 52961) to be used as controls. 48h after infection, cells were selected for three days with 2μg/ml puromycin (Sigma, P8833). To obtain pure knock-out clones, the pool of cells was single-cell grown, expanded, and expression of FBXW7 was analyzed by WB. The same procedure was followed for the generation of *ABCB1* ^-/-^ and *MCL1* ^-/-^ cells.

### CRISPR-Cas9 screens

For each screen, 5·10^6^ cells (50X library coverage) from previously described loss-of-function mES libraries^49^ were plated on gelatin. Cells were treated for approximately 10 days with the different compounds at doses in which no wild-type ES cells survive. For the UV-light screen, a single UV-light (254-nm UV-C) exposure was performed using a UVC 500 UV Crosslinker (Hoefer). Once there were less than 100 resistant clones, these were picked, isolated, and expanded. The resistance of individual clones was validated with the corresponding compound before following to the sequencing step. When the number of resistant clones exceeded 100, a pool of cells was grown, its resistance validated, and then the abundance of sgRNAs in the pool was quantified by PCR followed by next-generation sequencing. To identify the sgRNA sequences inserted in the single-isolated resistant clones, DNA was extracted and the fragment flanking the U6-sgRNA cassette was amplified by PCR and sequenced by Sanger sequencing. To identify the sgRNAs present in a pool of cells, DNA was extracted using a Gentra Puregene Blood Kit (Quiagen, 158445), following the manufacturer’s instructions. The U6-sgRNA cassette was then amplified by PCR using the KAPA HIFI Hot Start PCR kit (Roche, KK2502) and different tagged primers required for the subsequent Illumina sequencing. The PCR product was precipitated with sodium acetate 3M in EtOH 100% at −80°C for at least 20 min, pelleted and resuspended in water prior purification in agarose gel. Following a purity check of the PCR product, samples were sent for Illumina sequencing. sgRNA sequences were extracted from fastq files using Galaxy (https://usegalaxy.org/).

### Plasmids

The lentiviral plasmids pLentiCRISPR v2 (Addgene, 52961) and Lentiguide mCherry (kind gift of Cristina Mayor-Ruiz) were used to express sgRNAs in cells as described^79^. sgRNA sequences were designed using the MIT CRISPR design tool (http://www.genome-engineering.org/crispr/) and are available at **Table S1**. The lentiviral plasmid FUGW-eGFP (Addgene, 14883) was used to constitutively express eGFP. For RUBY3 expression, the eGFP sequence of the FUGW-eGFP vector was replaced with the RUBY3 cDNA^80^. The lentiviral plasmid pCLX-CHOP-dGFP (Addgene, 71299) was used to monitor CHOP transcription levels.

### Lentiviral production

Lentiviral vectors were individually co-transfected with third generation packaging vectors in HEK293T cells, using Lipofectamine 2000 (Invitrogen) to generate viral supernatants as previously described^81^. Lentiviral supernatants were collected 36h after transfection, pooled and passed through a 0.45 μM filter to eliminate cellular debris.

### RNA interference

Exponentially growing cells were trypsinised and transfected in suspension with 50nM of control siRNAs or human siRNAs targeting C-MYC (Horizon Discovery Biosciences, ON-TARGETplus siRNAs), following manufacturer’s instructions and using Lipofectamine RNAiMAX reagent (Thermo Fisher Scientific) and OPTIMEM medium (Life Technologies). For esiRNA libraries targeting mitochondrial factors (Sigma, MISSION^®^ esiRNA, **Table S2**), the same protocol was followed with 20nM of esiRNA and in a 96-well-plate format.

### Compounds

Compounds used in this study are indicated in **Table S3** and were used at the doses indicated in the Figure Legends. All compounds were dissolved in DMSO except cisplatin and oxaliplatin, which were dissolved in DMF; rapamycin and chloramphenicol, in EtOH; and doxycycline and minocycline, dissolved in sterile water. For the chemical screen we used a previously published in-house chemical library composed of 114 FDA-approved or in clinical trials anti-tumoral drugs solved in DMSO^82^. The library covers 80% of the pathways described in Reactome. The number of inhibitors for each pathway was the following: cell cycle (2), cell–cell communication (10), cellular response to external stimuli (11), chemotherapeutics (2), DNA repair (2), extracellular matrix reorganization (3), gene expression (12), hemostasis (16), immune system (20), metabolism proteins (2), organelle biogenesis and maintenance (1), and signal transduction (26).

### Flow Cytometry

For the analysis of mixed populations of cells expressing fluorescent proteins, cells at the corresponding mixture ratios were plated in 6-well tissue culture plates. The following day (or 8h after plating for mES), cells were treated with the indicated concentrations of drugs for 72h (unless specified), and then analysed by Flow Cytometry. For the analysis of CHOP transcription, DLD-1 cells expressing a CHOP-dGFP reporter cells were infected at 50% with Lentiguide mCherry vectors containing validated sgRNAs against the ISR kinases or an empty vector. 96h post-infection, cells were exposed to the indicated drugs for 72h. Cells were trypsinised, centrifuged, resuspended in PBS and incubated with DAPI for 10 minutes, and subsequently analysed the different cell populations using a flow cytometer BD FortessaTM (BD Biosciences).

For the analysis of mixed populations of cells expressing fluorescent proteins by High-Throughput flow cytometry, 4.000 cells were seeded in *μ*CLEAR bottom 96-well plates (Greiner Bio-One). The following day (or 8h for mES), cells were treated with the indicated concentrations of drugs for 72h (unless otherwise specified), and analysed by High-Throughput Flow Cytometry. For esiRNAs experiments, 4.000 cells were transfected in μCLEAR bottom 96-well plates (Greiner Bio-One). Every 3-4 days, a fraction of the culture was analysed by High-Throughput Flow Cytometry, while re-transfecting the rest with the esiRNAs. For all analyses, mixtures of cells were trypsinised, stained with DAPI, and the expression of the different fluorescent markers was analysed by using a BD FACS Canto II^TM^ (BD Biosciences) in High-Throughput mode. Data was processed with the Flow Jo 10^TM^ software to represent each cell population percentage.

### Viability assays

For clonogenic survival assays, 2.000 cells were plated in 6-well tissue culture plates in the corresponding culture medium. The following day, cells were treated with the indicated concentrations of drugs. Cells were maintained with the compounds for 10 days, changing the medium every 2-3 days, and then fixed and stained with 0.4% methylene blue in methanol for 30 min. Cell viability was also measured by High-Throughput Microscopy. In brief, 3.000 cells were seeded per well in μCLEAR bottom 96-well plates (Greiner Bio-One) and treated with the indicated concentrations of drugs the following day. 72h later, cells were fixed with 4% PFA and permeabilised with 0.5% Triton X-100, following standard procedures. Cells were subsequently stained with DAPI and images were automatically acquired from each well using an Opera High-Content Screening System (Perkin Elmer) or an ImageXpress Pico Automated Cell Imaging System (Molecular Devices). 20x or 10x magnification lenses were used indifferently, and images taken at non-saturating conditions. Images were then segmented using DAPI signals to generate masks that allowed the quantification of nuclei per condition.

### Immunofluorescence

For measuring ATF4 nuclear translocation, 8.000 cells were seeded per well in *μ*CLEAR bottom 96-well plates (Greiner Bio-One). The following day, cells were pre-treated for 1h with 50nM of ISRIB or DMSO and then treated with the indicated concentrations of drugs for 3 h. Next, cells were fixed with 4% PFA and permeabilised with 0.5% Triton X-100, following standard procedures. After blocking for 30 min, plates were stained with an anti-ATF4 primary antibody overnight (**Table S4**), followed by an anti-rabbit IgG-488 secondary antibody (Invitrogen, A21441) at 1:400 for 1h in RT on the following day. Plates were then stained with DAPI and images automatically acquired using an Opera High-Content Screening System (Perkin Elmer). A 20x magnification lens was used and images were taken at non-saturating conditions. Images were segmented using DAPI signals to generate masks matching cell nuclei, and the nuclear ATF4 intensity per cell was measured. For mitochondrial analyses, 8.000 cells were seeded per well in μ-slide 8-well plates (Ibidi). The following day, cells were fixed with 4% PFA and permeabilised with 0.5% Triton X-100, following standard procedures. After blocking for 30 min, cells were stained with an anti-citrate synthetase (CS) primary antibody for 30 min at 37°C (**Table S4**), and then with an anti-rabbit IgG-488 secondary antibody (Invitrogen, A21441) at 1:400 for another 30 min at 37°C. Cells were then stained with DAPI and images were acquired using a LEICA SP5 WLL confocal microscope. A 63x magnification lens was used and images were taken at non-saturating conditions. Images were segmented using DAPI and 488 signals to generate masks matching cell nuclei and mitochondria, respectively.

### Western Blotting

Cell pellets were lysed in 50mM Tris pH 7.9, 8M Urea and 1% Chaps followed by 30 min incubation with shaking at 4°C. For FBXW7 detection, pellets were lysed in 20mM HEPES pH 7.9, 0.4M NaCl, 1mM EDTA and protease inhibitors, followed by sonication and a 30 min incubation with shaking at 4°C. NuPAGE LDS (Life Technologies) with 10 mM DTT (Sigma) loading buffer was added to 20-30 μg of protein extracts, and samples were denatured for 10 min at 70 °C. For the detection of OXPHOS complexes, the denaturing step was performed at 50°C for 1h. Samples were run in precast gels and transferred for protein detection by using the corresponding primary antibodies (**Table S4**). The signal associated to HRP-conjugated secondary antibodies (ThermoFisher, mouse 31430 and rabbit 31460) was quantified using a SuperSignal™ West Pico PLUS Chemiluminescent Substrate kit (ThermoFisher, 34580) and a ChemiDoc MP Imagine System (BIO-RAD, 1708280).

### Mass spectrometry

Whole cell extract samples from WT and FBXW7-deficient DLD-1 or ES cells (2 biological replicates) were trypsin-digested using S-traps, isobaric-labelled with TMT^®^ 11-plex reagents, desalted using a Sep-Pak C18 cartridge and dried prior high pH reverse phase HPLC RP-HPLC pre-fractionation. Peptides were pre-fractionated offline by means of high pH reverse phase chromatography, using an Ultimate 3000 HPLC system equipped with a sample collector. Fractions were then analyzed by LC-MS/MS by coupling an UltiMate 3000 RSLCnano LC system to a Q Exactive Plus mass spectrometer (Thermo Fisher Scientific). Raw files were processed with MaxQuant (v1.6.0.16). Afterwards, the file was loaded in Prostar^83^ using the intensity values for further statistical analysis. Differential expression analysis was done using the empirical Bayes statistics limma. Proteins with a p-value < 0.05 and a log_2_ ratio higher than 0.27 (ES) or 0.3 (DLD-1) were defined as regulated, and the FDR was estimated to be below 2% by Pounds.

### RNA-seq

*FBXW7*^+/+^ and *FBXW7*^-/-^ DLD-1 cells were treated with DMSO or tigecycline (10*μ*M) for 24h. Total RNA was extracted from cell pellets using the Agilent Absolutely RNA Miniprep Kit following the manufacturer’s instructions. The sequencing library was constructed with the QuantSeq 3' mRNA-Seq Library Prep Kit (Lexogen), and approximately 10 million reads were obtained per sample by Illumina sequencing. Differential expression analyses were performed using Bluebee^®^ (Lexogen). For Gene Ontology (GO) analyses, the list of significantly upregulated genes (padj<0.1) was used as an input for the “The Gene Ontology Resource”, release 2021-09-01 (http://geneontology.org/)^84,85^. The transcriptional signature induced by tigecycline was also used as input at the Connectivity Map (CMap) Query clue.io tool (https://clue.io/)^65^ to identify drugs with signatures similar to that tigecycline.

### Animal studies

Athymic Nude-Foxn1nu 6-week female mice were acquired from Charles Rivers. 5·10^6^ exponentially-growing *FBXW7*^+/+^ and *FBXW7*^-/-^ DLD-1 cells were trypsinised and resuspended in PBS for injection in the flanks of 8-week mice. 6 days after, mice were randomized into three groups per genotype (six groups in total, 10 mice per group) and treatment was started with 1,5 mg/kg paclitaxel (*in vivo* reference in **Table S3**), 50 mg/kg of tigecycline (*in vivo* reference in **Table S3**) or vehicle via intraperitoneal (i.p.) injection, on a three times per week schedule. Tumours were measured every 2-3 days, and once they reached 1600 mm^3^ (measures were calculated using the standard formula length x width x 0.5), mice were sacrificed and their tumours extracted. Health status of mice was monitored daily. Mice were maintained under standard housing conditions with free access to chow diet and water, as recommended by the Federation of European Laboratory Animal Science Association. All mice work was performed in accordance with the Guidelines for Humane Endpoints for Animals Used in Biomedical Research, and under the supervision of the Ethics Committee for Animal Research of the “Instituto de Salud Carlos III”.

### Bioinformatic analyses

#### Drug resistance

Drug responses associated with wild type and *FBXW7*^mut^ cancer cell lines from the NCI-60 and the CCLE collections were extracted and downloaded from the Genomics and Drugs integrated Analysis (GDA) (http://gda.unimore.it/) portal. The same analysis was performed for wild type and *ABCB1*^mut^ cell lines of the NCI60 (there was no data associated with ABCB1 mutations available in the CCLE). The linear model analysis between *FBXW7* expression levels and the Area Under the Curve (AUC) associated with drugs was performed using the CTRP (https://portals.broadinstitute.org/ctrp.v2.1/) portal. The plot represents the coefficient of the compounds which reached statistical significance. A R version compatible with version 3.6.3 was employed for analysis and representation of data.

#### GDC Pan-Cancer survival data

The UCSC XenaBrowser (https://xenabrowser.net/) was used to explore the GDC Pan-Cancer database containing data on patients’ survival, drug treatment and tumour *FBXW7* gene expression. The data was downloaded, and separated into two datasets: one containing data from patients under treatment and the other from patients without drug treatment information. For each of these datasets, patient’s data was stratified into two groups according to the median of *FBXW7* expression, and survival data from each group was plotted. A Cox regression, including tumour type as co-variable, was also performed. R version compatible with version 3.6.3 was employed for analysis and representation of data.

#### CCLE proteomics

For proteomic analyses of CCLE, data was extracted from a recently published work^86^. 388 cancer cell lines were classified according to *FBXW7* mutational and copy number variation status. Only cell lines harbouring coding, damaging, or non-conserving alterations in *FBXW7* were labelled as mutated. Cell lines with an absolute copy number score of 0 for *FBXW7* were also included. Differential expression analysis was carried out using limma^87^ on normalized expression levels between *FBXW7-mutant* and WT cancer cell lines. For analysis and representation of the data, R version 3.6.1 was used. For GSEA analyses of proteomics data, GSEA v2.2.4. was used, using a list of pre-ranked Fold Change values as an input.

### Statistics and reproducibility

No statistical method was used to predetermine sample size. Experiments using cells and mice were randomized. Most experiments were performed three times with two technical replicates per experiment, in two independent mutant clones and either the mean of the three experiments or a representative example is shown in the Figures. Values are reported as the mean ± SD, except for the animal studies, were the mean ± SEM is indicated. No data were excluded from the analyses. For general assessment of the difference between two sets of data, we used the two-tailed unpaired Student’s t-test (GraphPad Prism version 7.04). We tested for statistical significance of the overlap between the two groups of genes using the normal approximation of the exact hypergeometric test (http://nemates.org/MA/progs/overlap_stats.cgi). The computational methods used for the differential expression analyses, GO analyses, GSEA studies and other bioinformatics analyses, are explained in their specific methods section. n. Unless otherwise indicated, threshold FDR and padj values were set at <0.05. P-values <0.05 were considered to be significant (n.s. p>0.05, *p<0.05, **p<0.01, ***p<0.001). All statistical parameters, tests and other relevant detailed information are reported in the Figures and corresponding Figure Legends.

### Data availability

Correspondence and request of materials should be addressed to O.F. RNA sequencing data associated with this work are accessible at the GEO repository, under accession number GSE189499. Mass spectrometry proteomic datasets are available at the PRIDE repository with accession numbers PXD029981.

