## Supplementary material for "Activation of the Integrated Stress Response overcomes multidrug resistance in FBXW7-deficient cells": Figures S1-8

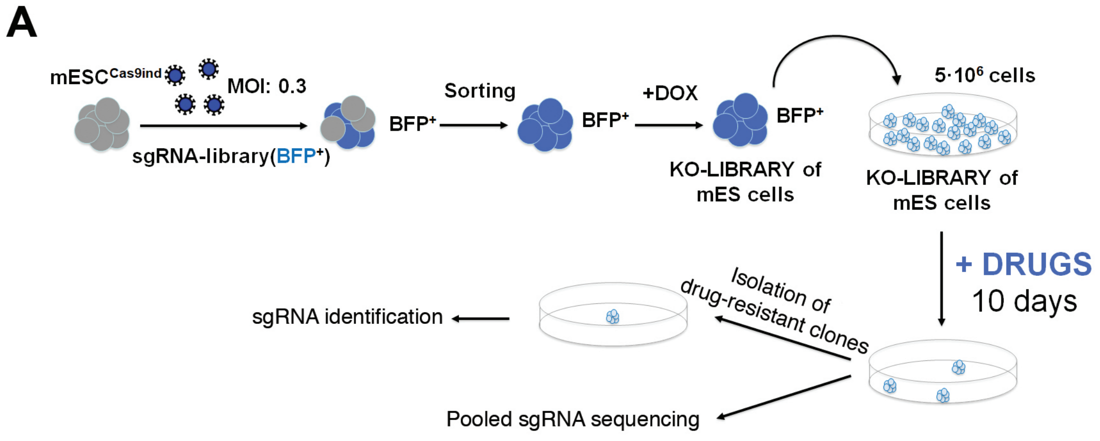


**B**

Drug-resistant clones with sgRNAs targeting *Fbxw7*

| | Library 1 | Library 2 |
|---|---|---|
| CISPLATIN | 8 / 11 | 1 / 11 |
| DAB-III | 9 / 15 | - |
| RIGOSERTIB | 5 / 8 | - |
| CSCi | - | 5 / 20 |
| UV | - | 5 / 25 |

**C**

Pooled sgRNA sequencing

| | sgRNA sequence | Pos. | Nº reads |
|---|---|---|---|
| CISPLATIN LIBRARY 1 | *Fbxw7* sgRNA (1) | 1 | 362449 |
| CISPLATIN LIBRARY 2 | *Fbxw7* sgRNA (1) | 20 | 1465 |
| | *Fbxw7* sgRNA (2) | 44 | 178 |
| UV LIBRARY 1 | *Fbxw7* sgRNA (1) | 7 | 38937 |
| | *Fbxw7* sgRNA (2) | 32 | 5733 |
| | *Fbxw7* sgRNA (3) | 9 | 35081 |
| | *Fbxw7* sgRNA (4) | 204 | 340 |

**Fig. S1. Enrichment of *FBXW7*-targeting sgRNAs in CRISPR screens.** (**A**) Pipeline of CRISPR-Cas9 screens. Briefly, mES carrying a doxycycline-inducible Cas9 were infected at a low multiplicity of infection (MOI 0.3) with a lentiviral library of sgRNAs targeting virtually all mouse genes. Infected cells were sorted by FACS based on BFP expression, after which doxycycline was added for 10 days. Mutagenized mES libraries were then isolated and used for the screens. $5 \cdot 10^6$ cells were used per screen (50X library coverage) and exposed for around 10 days to test different compounds at doses that kill all the WT mES cells. Treatment resistant clones, if any, were then isolated and expanded, and sgRNA sequences amplified by PCR and identified by Sanger sequencing. If there were more than 100 resistant clones, they were pooled together and sgRNAs were identified by Illumina sequencing after amplification of the sgRNAs by PCR. (**B**) Number of clones carrying *Fbxw7*-targeting sgRNAs in drug resistant clones isolated during genetic screens against the indicated drugs. The screens were conducted in 2 different libraries coming from 2 independent mES clones. (**C**) *Fbxw7*-targeting sgRNAs identified by Illumina sequencing in the resistant pool of cells coming the genetic screens against UV and cisplatin. The numbers inside parentheses indicate different sgRNA sequences. The position in the enrichment-rank of the screen and the number of reads per sgRNA is also indicated.

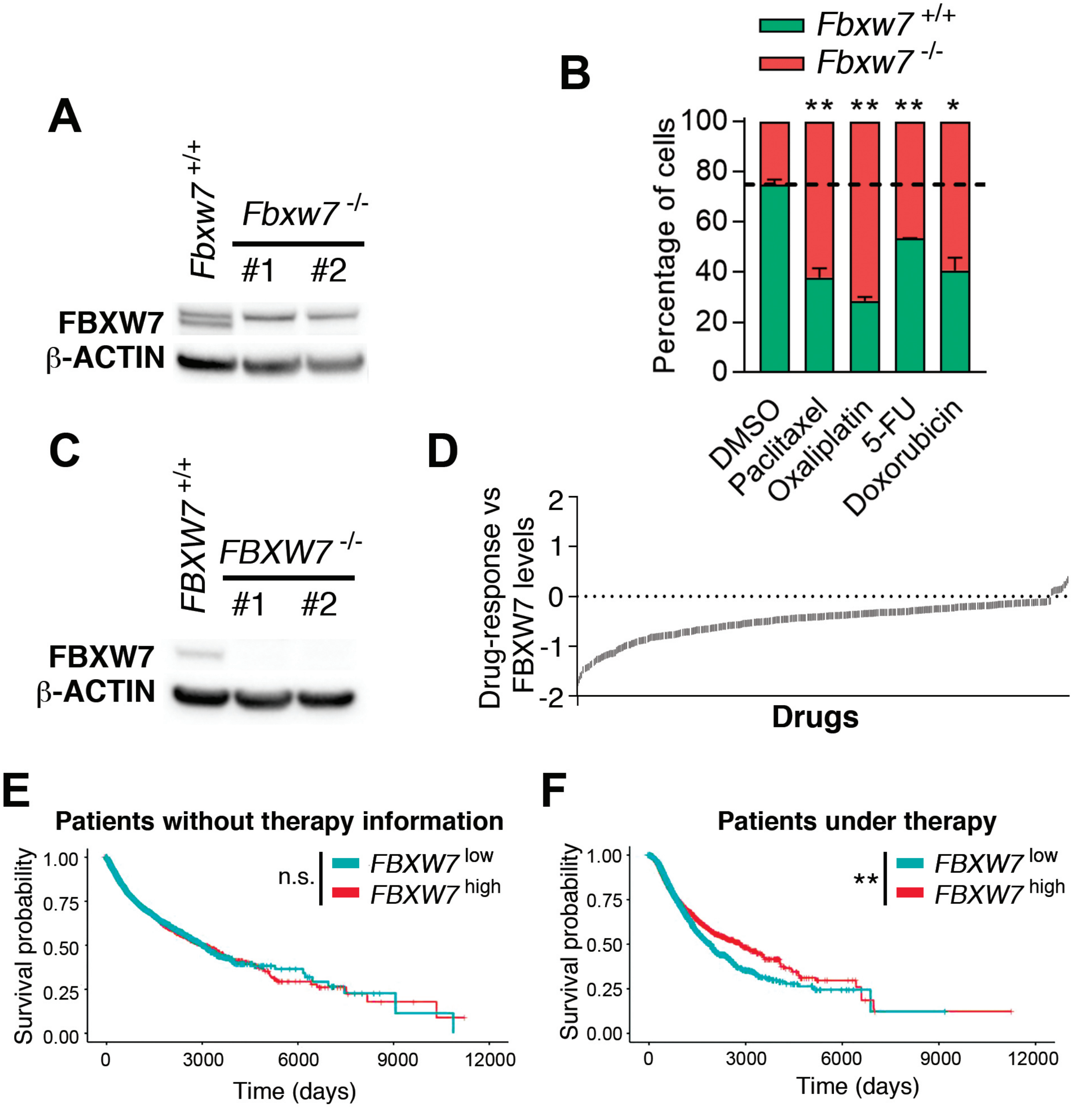


**Fig. S2. FBXW7 deficiency leads to MDR.** (**A**) WB illustrating the absence of FBXW7 expression in 2 independent *Fbxw7* deficient mES clones generated by CRISPR editing. β-ACTIN levels are shown as a loading control. (**B**) Percentage of viable *Fbxw7*$^{+/+}$ and *Fbxw7*$^{-/-}$ mES cells 48h after being treated with paclitaxel (30nM), oxaliplatin (750nM), 5-FU (2µM) and doxorubicin (25nM). The culture started with a 1:3 ratio of *Fbxw7*$^{+/+}$ and *Fbxw7*$^{-/-}$ cells. The experiment was repeated three times, and a representative example is shown. Error bars indicate SD. n.s. $p>0.05$, *$p<0.05$, **$p<0.01$, ***$p<0.001$ (t-test). Cell percentages were quantified by flow cytometry. (**C**) WB illustrating the absence of FBXW7 expression in 2 independent FBXW7-deficient DLD-1 clones generated by CRISPR editing. β-ACTIN levels are shown as a loading control. (**D**) Representation of the coefficients resulting from a lineal model analysis between FBXW7 expression and the AUC of multiple therapeutic compounds in cell lines of the CTRP dataset. Each line represents a compound. Negative values indicate resistance to the compound. (**E,F**) Survival probability in cancer patients for which there is no treatment information (n=15156 number of patients) (**E**) and in those under any therapy (n=5001 number of patients) (**F**), stratified by *FBXW7* mRNA levels (above or below median values). Data comes from the GDC Pan-Cancer study. n.s. $p>0.05$, *$p<0.05$, **$p<0.01$.

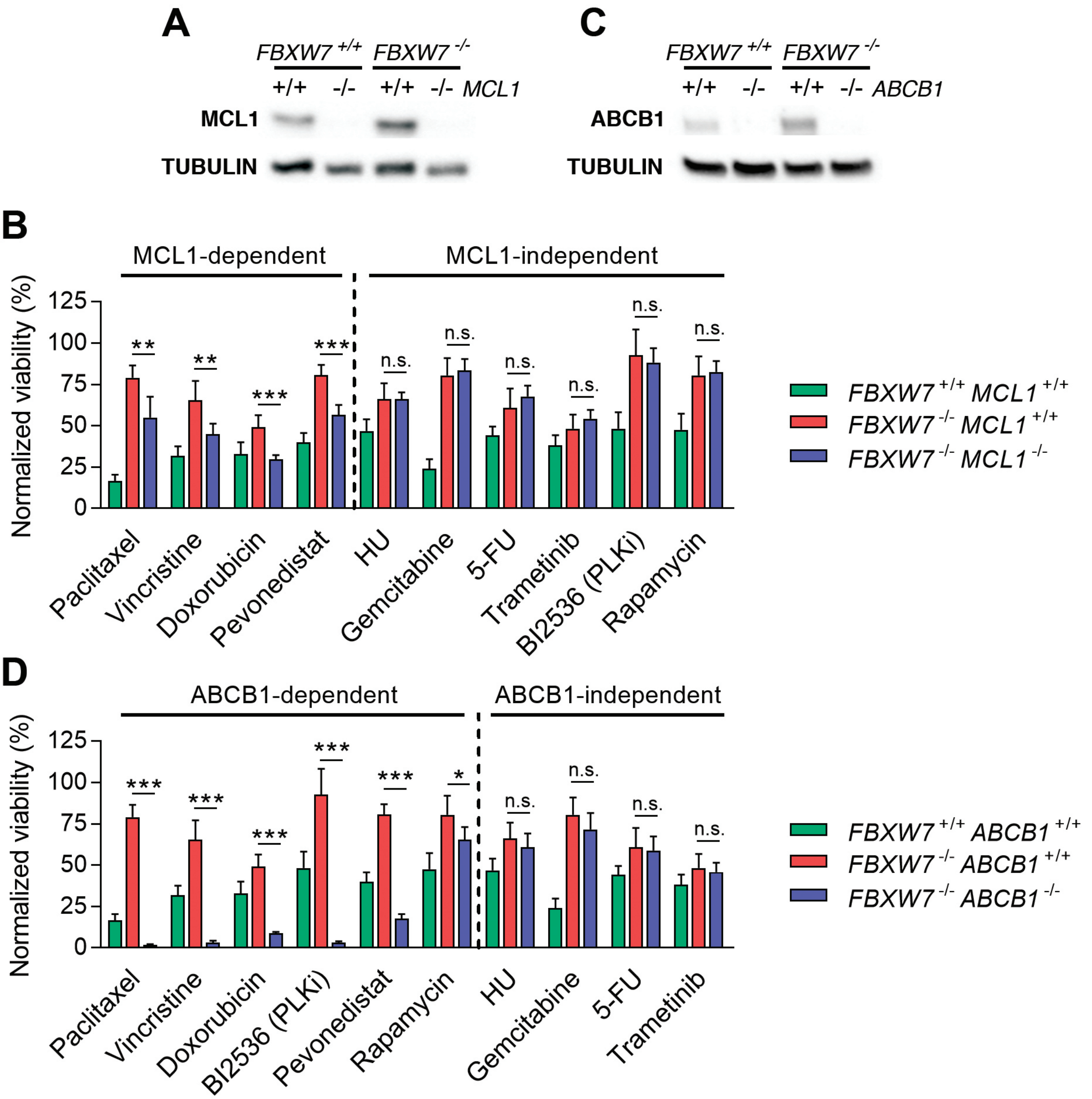


**Fig. S3. Impact of *MCL1* or *ABCB1* deletion on the resistance associated to FBXW7 deficiency.** (**A,B**) WB illustrating the absence of MCL1 (**A**) and ABCB1 (**B**) expression in *FBXW7*$^{+/+}$ and *FBXW7*$^{-/-}$ DLD-1 cells generated by CRISPR editing. TUBULIN levels are shown as a loading control. (**C**) Normalized viability of *FBXW7*$^{+/+}$*MCL1*$^{+/+}$, *FBXW7*$^{-/-}$*MCL1*$^{+/+}$ and *FBXW7*$^{+/+}$*MCL1*$^{-/-}$ DLD-1 cells after treatment with paclitaxel (40nM), vincristine (10nM), doxorubicin (25nM), hydroxyurea (HU, 75µM), gemcitabine (10nM), Fluorouracil (5-FU, 10µM), trametinib (5µM), BI2536 (PLK1i, 10nM), pevonedistat (200nM) and rapamycin (10µM) for 72h. Cell nuclei were quantified by high-throughput microscopy (HTM) upon staining with DAPI. Equivalent results were seen with an independent MCL1-deficient clone. Error bars indicate SD (n=3). n.s. p>0.05, **p<0.01, ***p<0.001 (t-test). (**D**) Normalized viability of *FBXW7*$^{+/+}$*ABCB1*$^{+/+}$, *FBXW7*$^{-/-}$ *ABCB1*$^{+/+}$ and *FBXW7*$^{+/+}$ *ABCB1*$^{-/-}$ DLD-1 cells after treatment with the same drugs and doses indicated in (**C**) for 72h. Cell nuclei were quantified by high-throughput microscopy (HTM) upon staining with DAPI. The experiment was repeated three times, and a representative example is shown. Equivalent results were seen with an independent ABCB1-deficient clone. Error bars indicate SD (n=3). n.s. p>0.05, **p<0.01, ***p<0.001 (t-test).

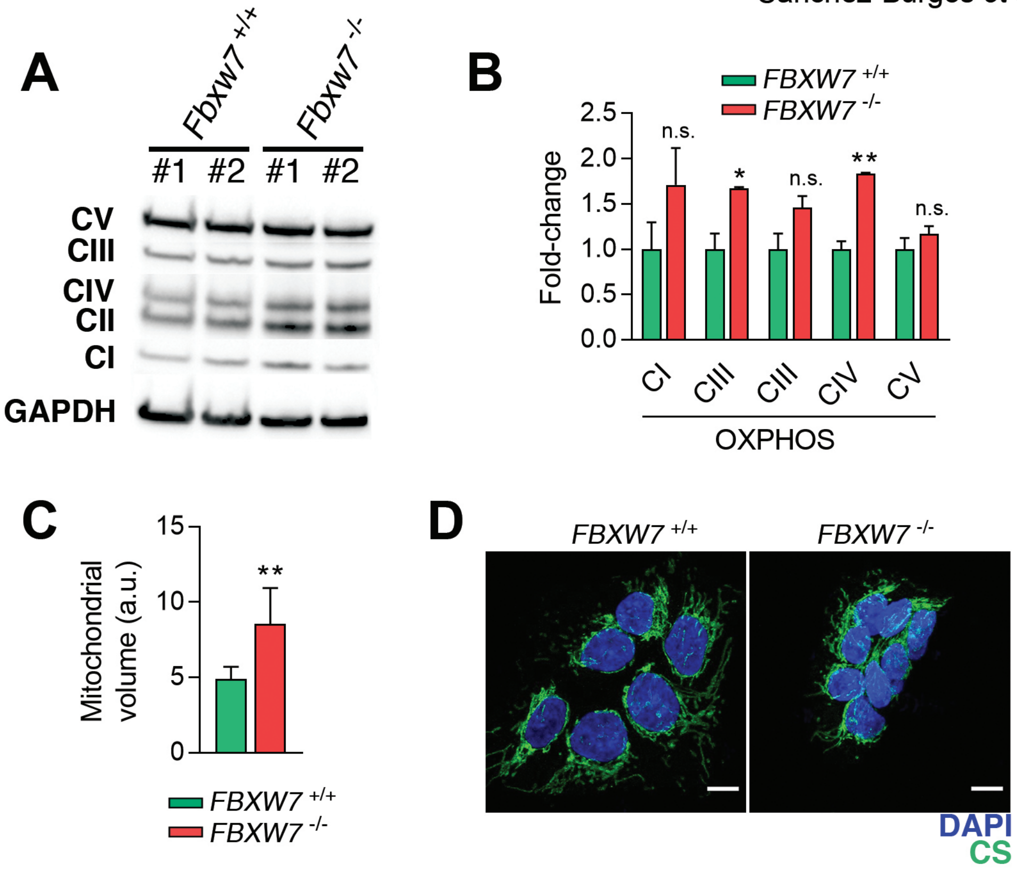


**Fig. S4. FBXW7 deficiency is associated to an increased mitochondrial volume.** (**A**) WB illustrating the levels of the different mitochondrial OXPHOS complexes in 2 independent clones of *Fbxw7*$^{+/+}$ and *Fbxw7*$^{-/-}$ mES cells. (**B**) Quantification of the data from (**A**). Error bars indicate SD. n.s. $p>0.05$, $*p<0.05**p<0.01$ (t-test). (**C**) Mitochondrial volume in *FBXW7*$^{+/+}$ and *FBXW7*$^{-/-}$ DLD-1 cells as quantified from the levels of the mitochondrial factor citrate synthase (CS) by HTM. DAPI staining was used to stain nuclei. Error bars indicate SD. $**p<0.01$ (t-test). This experiment was repeated three times and a representative example is shown. (**D**) Representative images from (**E**). Scale bar (white) indicates 10µm.

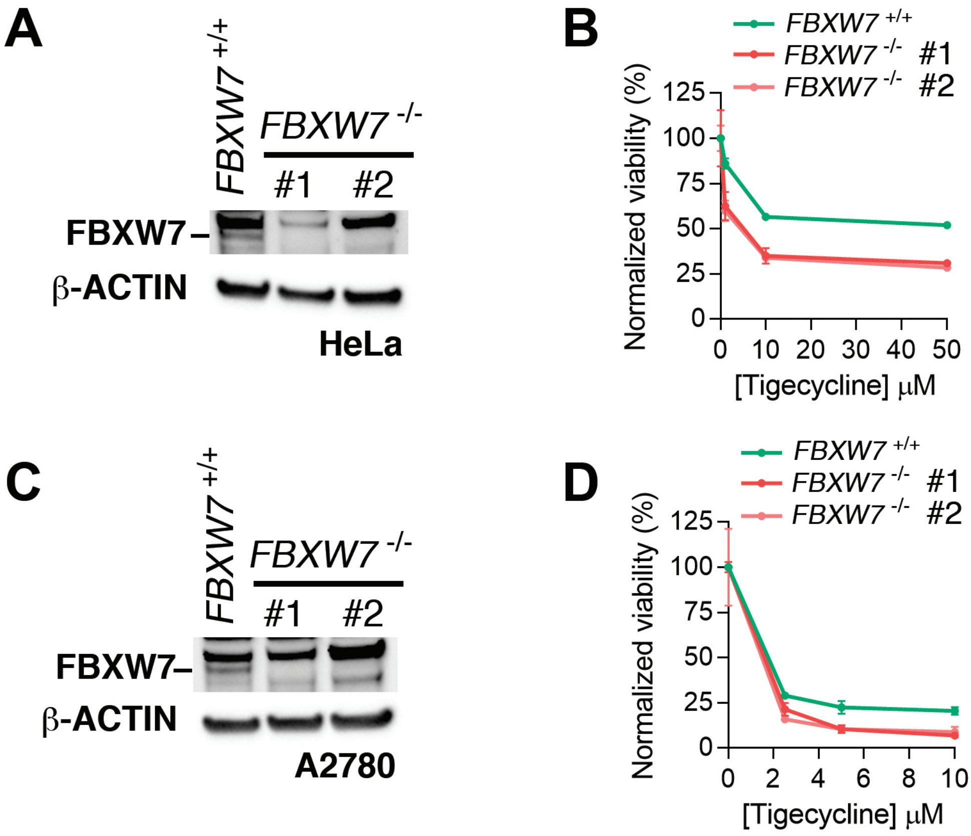


**Fig. S5. Sensitivity to tigecycline in FBXW7-deficient cell lines.** (**A**) WB illustrating the absence of FBXW7 expression in 2 independent FBXW7-deficient clones generated by CRISPR editing in HeLa cells. β-ACTIN levels are shown as a loading control. (**B**) Normalized viability of *FBXW7*$^{+/+}$ and 2 independent clones of *FBXW7*$^{-/-}$ HeLa cells upon treatment with increasing doses of tigecycline for 72h. Cell nuclei were quantified by high-throughput microscopy (HTM) upon staining with DAPI. The experiment was repeated three times, and a representative example is shown. Errors indicate SD. (**C**) WB illustrating the absence of FBXW7 expression in 2 independent *FBXW7* deficient clones generated by CRISPR editing in A2780 cells. β-ACTIN levels are shown as a loading control. (**D**) Normalized viability of *FBXW7*$^{+/+}$ and 2 independent clones of *FBXW7*$^{-/-}$ A2780 cells upon treatment with increasing doses of tigecycline for 72h. Cell nuclei were quantified by high-throughput microscopy (HTM) upon staining with DAPI. The experiment was repeated three times, and a representative example is shown. Errors indicate SD.

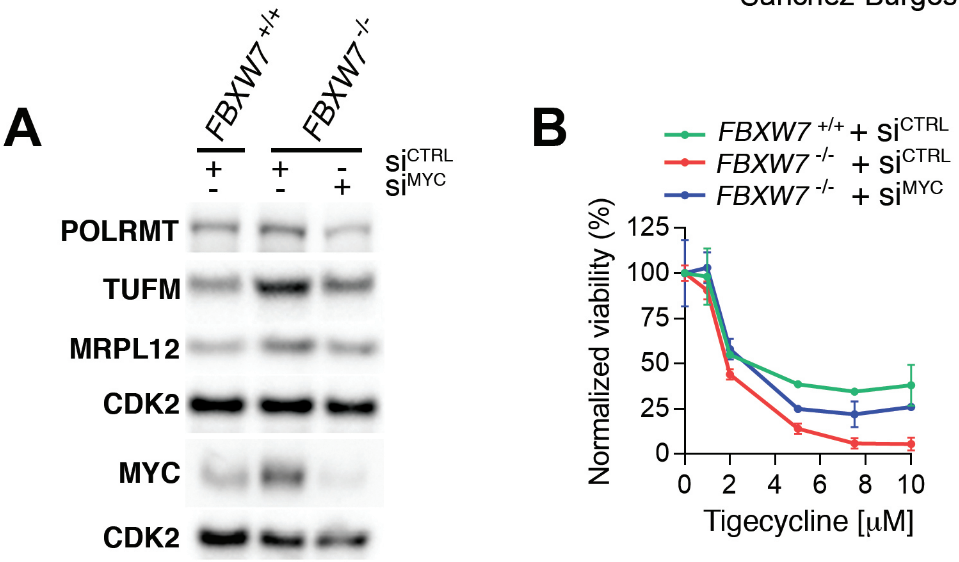


**Fig. S6. MYC depletion reduces the expression of mitochondrial factors and the sensitivity to tigecycline of FBXW7-deficient cells.** (**A**) WB illustrating the levels of the mitochondrial factors POLRMT, TUFM and MRPL12 in *FBXW7*$^{+/+}$ and *FBXW7*$^{-/-}$ DLD-1 cells 48h after transfection with siRNAs targeting MYC or a control siRNA. MYC levels are also shown, which were evaluated in an independent WB. CDK2 was used as a loading control in both WB. (**B**) Normalized viability of *FBXW7*$^{+/+}$ and *FBXW7*$^{-/-}$ DLD-1 cells transfected with siRNAs targeting MYC or a control siRNA upon treatment with increasing doses of tigecycline. Cell nuclei were quantified by high-throughput microscopy (HTM) upon staining with DAPI. The experiment was repeated three times, and a representative example is shown. Errors indicate SD.

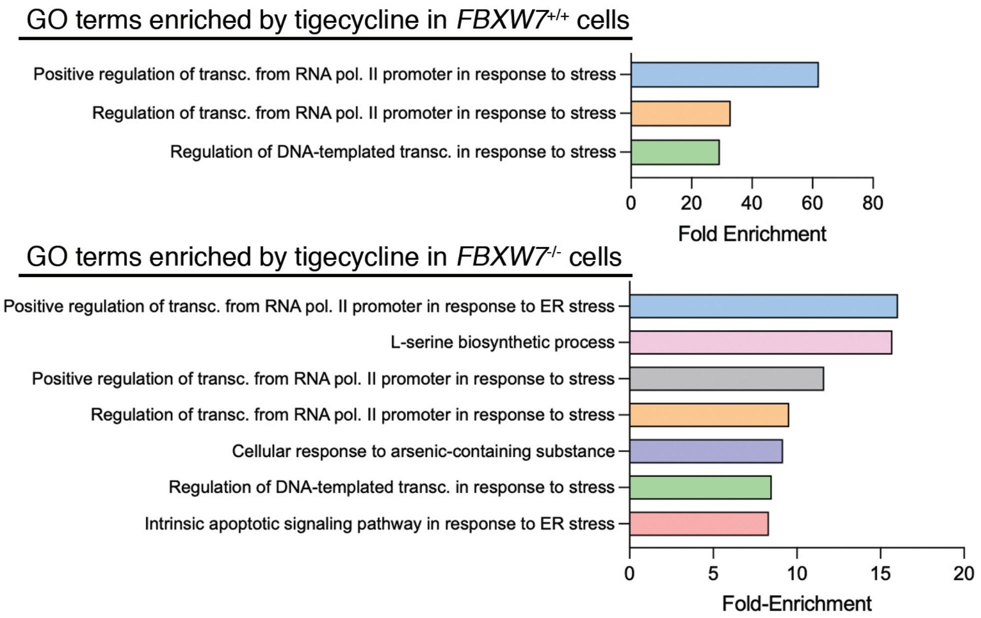


**Fig. S7. Transcriptional effects of tigecycline in *FBXW7*$^{+/+}$ and *FBXW7*$^{-/-}$ DLD-1 cells.** Most significantly enriched Gene Ontology (GO) terms identified in RNAseq analyses of *FBXW7*$^{+/+}$ (top) and *FBXW7*$^{-/-}$ (bottom) DLD-1 cells treated with tigecycline (10 μM) for 24h.

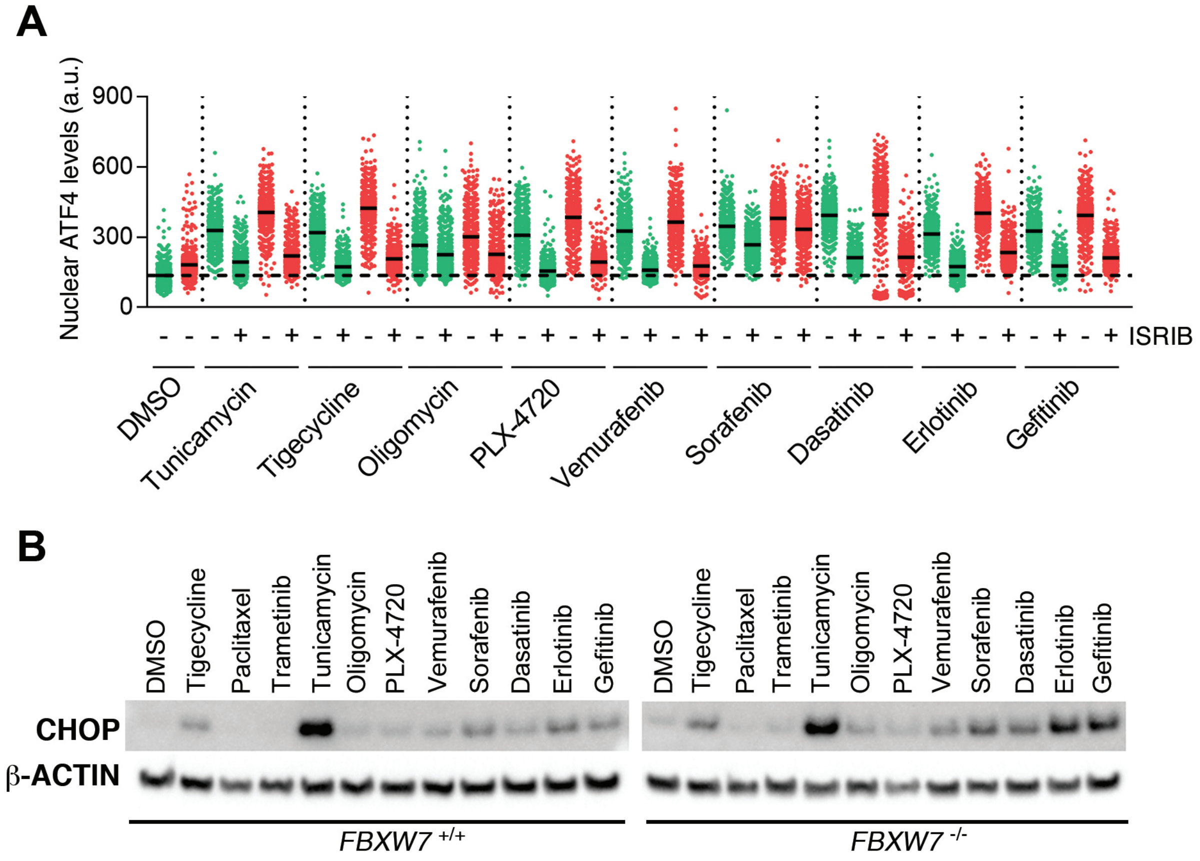


**Fig. S8. Activation of the ISR by drugs that overcome the MDR of FBXW7-deficient cells.** (**A**) Nuclear ATF4 levels quantified by HTM in DLD-1 cells upon treatment with 10 μM of the indicated compounds (except tunicamycin, which was used at 1μM) with or without the ISR inhibitor ISRIB (50nM) for 3h. This experiment was performed 3 times, and a representative example is shown. (**D**) WB illustrating the levels of CHOP in *FBXW7* $^{+/+}$ and *FBXW7* $^{-/-}$ DLD-1 cells treated as in (**A**). Paclitaxel and trametinib were also added at 250 nM and 10 μM, respectively. β-ACTIN levels are shown as a loading control.
